# Synthetic spider silk forming highly-aligned nanoarchitectures on 2D-material surfaces

**DOI:** 10.1101/2024.01.27.577543

**Authors:** Dominic R Whittall, Ryo Kato, Akari Okimura, Rio Takizawa, Kazunori Motai, Hiroki Maeda, Yui Yamazaki, Taka-aki Yano, Eriko Takano, Yuhei Hayamizu

## Abstract

The synergy between silk proteins and nanomaterials can lead to novel materials with improved mechanical and electrical properties. Designed peptides have been previously utilised in the functionalisation of two-dimensional material surfaces in a self-assembly manner, including graphite, to develop highly sensitive electrical biosensors. These studies have predominantly focused on functionalising the surfaces with peptides of less than several kDa in size. In this work, we assessed the capability of a ∼35 kDa synthetic spider silk protein to serve as a candidate biomolecular scaffold on a range of two-dimensional materials: graphite, molybdenum disulphide and boron nitride. The structural properties of the synthetic spider silk protein at the 2D material surface were characterised at the nanoscale for the first time using a multi-analysis approach incorporating atomic force microscopy and fluorescence microscopy, in addition to polarised Raman and tip-enhanced Raman spectroscopy. The synthetic spider silk protein was revealed to self-assemble into stable nanowire structures of monolayer thickness. Our findings have demonstrated the feasibility of functionalising 2D materials with spider silk-based proteins and will unlock new possibilities in the development of next-generation high-performance biosensing devices.

## Introduction

Silk proteins have been intensively studied due to their outstanding properties in terms of mechanical strength, biocompatibility and biodegradability. Silk proteins have immense potential for application within a diverse range of fields, including tissue engineering, drug delivery and next-generation textiles [1-3]. The combination of silk proteins and nanomaterials can lead to novel biomaterials displaying enhanced mechanical and electrical properties. The resulting composites have potential applications in areas such as biomedical engineering, energy storage and bioelectronics. Nanocarbons such as nanotubes and graphene have been widely used for composites [4-7], and more recently other two-dimensional (2D) materials have also attracted interest [8,9]. The atomically flat surface of graphene and other 2D materials, such as molybdenum disulphide (MoS_2_) and hexagonal boron nitride (h-BN), is a versatile platform for engineering the interface between nanomaterials and biomaterials [10-14].

Of all the potential applications, electrical biosensing with 2D materials is one of the most promising. For example, graphene field-effect transistors (GEFTs) are used for the detection of target molecules in solution, in which the graphene surface is functionalised to enable specific binding affinities towards the target molecules. Peptides mimicking the amino acid sequences of silk proteins have demonstrated their ability to functionalise the graphene surface [15], and several studies have reported successful biosensing [1, 16-18]. Although peptides are useful for biosensing, there remains a considerable gap in molecular size between peptides and functional proteins that can be used as scaffolds for effective biosensing. Typical peptides used in graphene functionalisation are around 1 kDa, much smaller than many other reliable functional proteins. Moreover, to maintain electrical sensitivity, scaffold thickness must be less than the Debye length of >2 nm for use in a standard physiological solution or buffer [18]. Therefore, in order to develop versatile molecular scaffolds for use in electrical biosensing, it is necessary to identify novel functional proteins with stable, highly-ordered structural conformations. In addition, these proteins should be capable of adopting uniform morphologies at the material surface.

Among the silk proteins, spider silk proteins arguably hold the greatest potential. A highly versatile biomaterial, fibres of major ampullate (dragline) silk possess a tensile strength greater than steel and toughness in excess of various synthetic fibres [19, 20]. In addition, the biocompatibility of dragline silk makes it attractive for a variety of uses in the field of biomedicine, including scaffolds for tissue regeneration, implant coatings and vehicles for drug delivery [2]. However, native silks are difficult to harvest from their natural producers, and both the size (250–350 kDa) and repetitive amino acid composition render heterologous expression of native spider silk proteins, known as spidroins, problematic [21], Consequently, smaller recombinant spidroins (mini-spidroins) that retain many of the desirable properties of their native counterpart are highly attractive for certain practical applications [22-24] (Figure 1).

**Figure 1:**
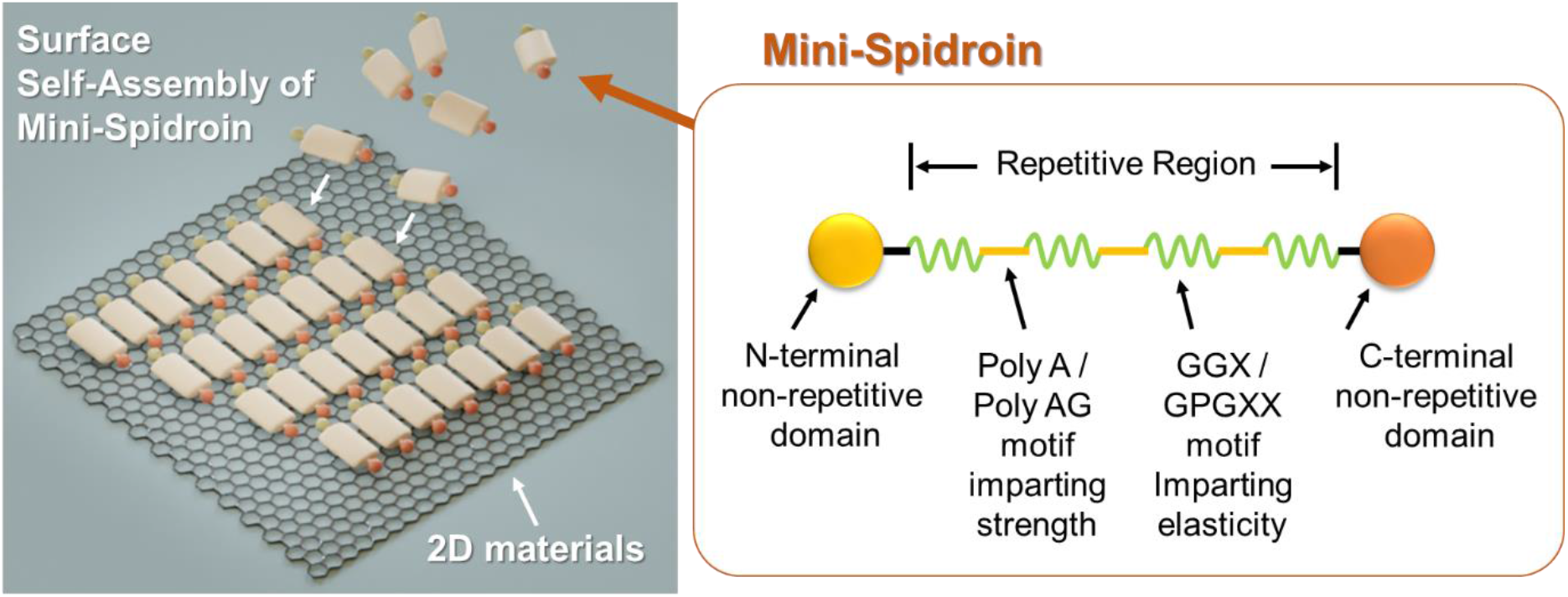
Hypothesised general mechanism of mini-spidroin assembly at a 2D material surface and the modular structure of mini-spidroin protein. The non-repeating N-terminal and C-terminal domains flank a larger repetitive section, which include alternating polyA or polyAG and GGX or GPGXX motifs.

Native spidroins form stable inter- and intra-β-sheet structures via a dense network of hydrogen bonds [25]. These β-sheet nanocrystals, present within a semi-amorphous protein matrix of α-helices and β-turns, underpin the outstanding strength of spider silk fibres [26]. The hierarchical structure of spider silk has been extensively studied using a range of high-resolution spectroscopic techniques, including scanning electron microscopy [27, 28] atomic force microscopy [29, 30] and transmission electron microscopy [28, 31, 32]. The highly-ordered, conformationally stable secondary structure of spidroins suggest their suitability for use as molecular scaffolds.

In this work, we investigated the potential of a ∼35 kDa recombinant spider silk protein (NR7C) as a new candidate molecular scaffold on graphite and other 2D materials for use in electrical biosensing. Herein, we report the surface self-assembly of recombinant mini-spidroin NR7C across three 2D material surfaces: graphite, MoS_2_ and h-BN. The morphology of the self-assembled structures of mini-spidroin NR7C on each substrate was characterised using atomic force microscopy (AFM). The conformational properties of the mini-spidroin crystal were investigated by polarised Raman spectroscopy. Further, the conformational properties of the self-assembled mini-spidroin structures on single-layer MoS_2_ were characterised at the single protein level using tip-enhanced Raman spectroscopy. The present demonstration of self-assembly of recombinant mini-spidroin on nanomaterials together with morphological and chemical characterisation of their properties would pave the way towards functionalisation of the surface of nanomaterials, thereby accelerating the development of next-generation high-performance biosensing devices.

## Results and Discussion

### Characterisation of self-assembled structures by Atomic Force Microscopy

In this work, three different 2D materials: Graphite, MoS_2_, and h-BN were utilized as scaffold for spidroin NR7C. Self-assembly of purified recombinant spidroin NR7C on each 2D material was examined at various concentrations of proteins in the solution. The protein solutions with concentrations of 0.5, 2.0, and 10 ug/ml were placed on each 2D material to investigate the self-assembly on the surfaces. Immediately following protein incubation, the samples were thoroughly rinsed with DI water and then dried with nitrogen gas. Their morphology on the surface was characterised by AFM (Figure 2).

**Figure 2:**
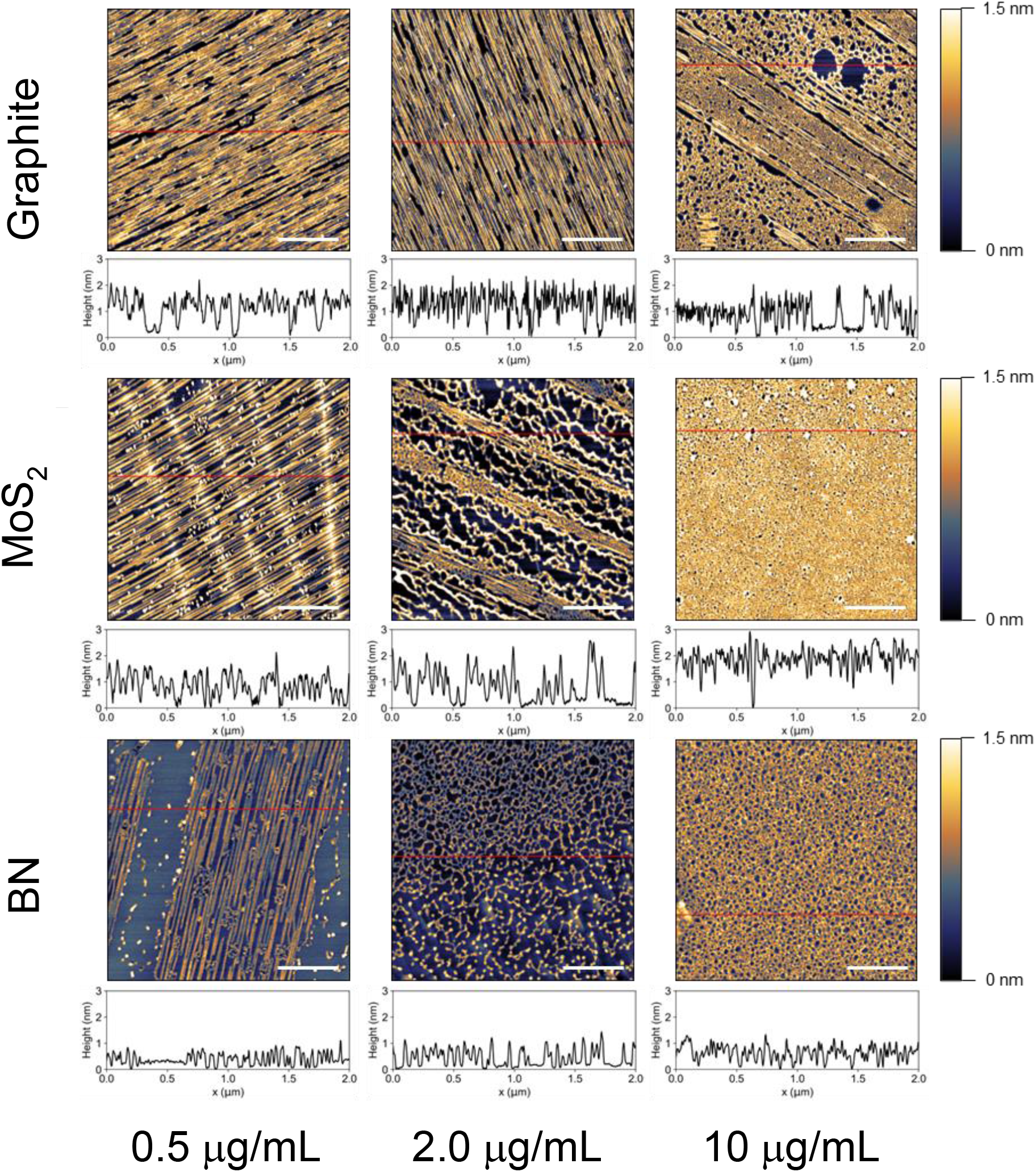
AFM images of NR7C spidroin self-assembly on 2D-material substrates (Graphite; Molybdenum disulphide; Boron nitride). Spidroin morphology, height and coverage are dependent on the concentration of NR7C and the type of 2D-material material substrate. Solutions of NR7C were prepared in DI water to concentrations of 0.5, 2 and 10 μg/mL. The scale bar is 500 nm.

Our study found that NR7C binds to the surface with high coverage across a broad range of concentrations (0.5 – 10 ug/ml). At the lowest examined concentration of 0.5 ug/ml, NR7C forms uniform nanowires on each of the 2D materials studied. The direction of the nanowires on each material surface was aligned, spanning lengths in excess of 2 μm. The ability of NR7C, a relatively large protein of ∼35 kDa, to form long-range, ordered structures indicates the presence of strong inter- and intramolecular interactions between and amongst monomers of NR7C at the surface.

The thickness of the NR7C nanowires formed on both graphite and MoS_2_ surfaces was approximately 2 nm. Interestingly, nanowires formed on h-BN were thinner than those formed on the other surfaces and were between 0.5 and 1 nm in thickness. Nanowires with a thickness of ∼0.5 nm corresponded to the monomolecular layer of NR7C at the h-BN surface. Consequently, the two-fold difference in nanowire thickness implies the formation of bilayer NR7C structures of NR7C at the h-BN surface.

Nanowire morphology, in addition to the propensity of NR7C to adopt nanowire or amorphous structures on surfaces, appeared highly dependent on protein concentration. At the highest examined concentration of 10 ug/ml, linear nanowire structures either partially (graphite) or totally (MoS_2_ and h-BN) disappeared. The amorphous structures adopted by NR7C suggest that a high-density protein concentration provides fewer opportunities for NR7C to diffuse and interact with its adjacent monomers, hampering the formation of well-aligned, ordered nanowires. The transition state between aligned nanowires and amorphous structures was visible at the middle concentration of 2 ug/ml on both MoS_2_ and h-BN surfaces. As the transition state appears to occur at 10 ug/ml on the graphite surface, the threshold concentration at which the transition state can be observed appears to be different on each substrate. This suggests that the interaction between NR7C monomers and the surface are unique to the substrate material in question. Graphite, MoS_2_, and h-BN surfaces are highly hydrophobic [33-35]. In addition, NR7C is also both hydrophobic (GRAVY index = 0.051) and stable (Instability index = 32.74) [36]. Hydrophobic surfaces have a stronger binding affinity for proteins compared to hydrophilic surfaces. This is due to the increased interactions between the hydrophobic surface and the hydrophobic domains of the protein that are exposed during adsorption. Hydrophobic surfaces displace interfacial water more easily, resulting in a stronger binding compared to hydrophilic surfaces with stronger water binding [37]. Hence, the presence of ordered NR7C nanostructures following rinsing with DI water further suggests the formation of highly stable β-sheet structures at the 2D material surfaces.

### Fluorescence Microscopy

NR7C nanostructures formed at the surface of h-BN flakes were further characterised by fluorescence microscopy (Figure 3). This fluorescent imaging allows us to identify the domain size of the self-assembled nanowires of NR7C in the larger field of view compared with AFM. We employed Thioflavin T (ThT) as a fluorescent label to stain the nanowires of NR7C to visualize their macro structures after the self-assembly on the surface.

**Figure 3:**
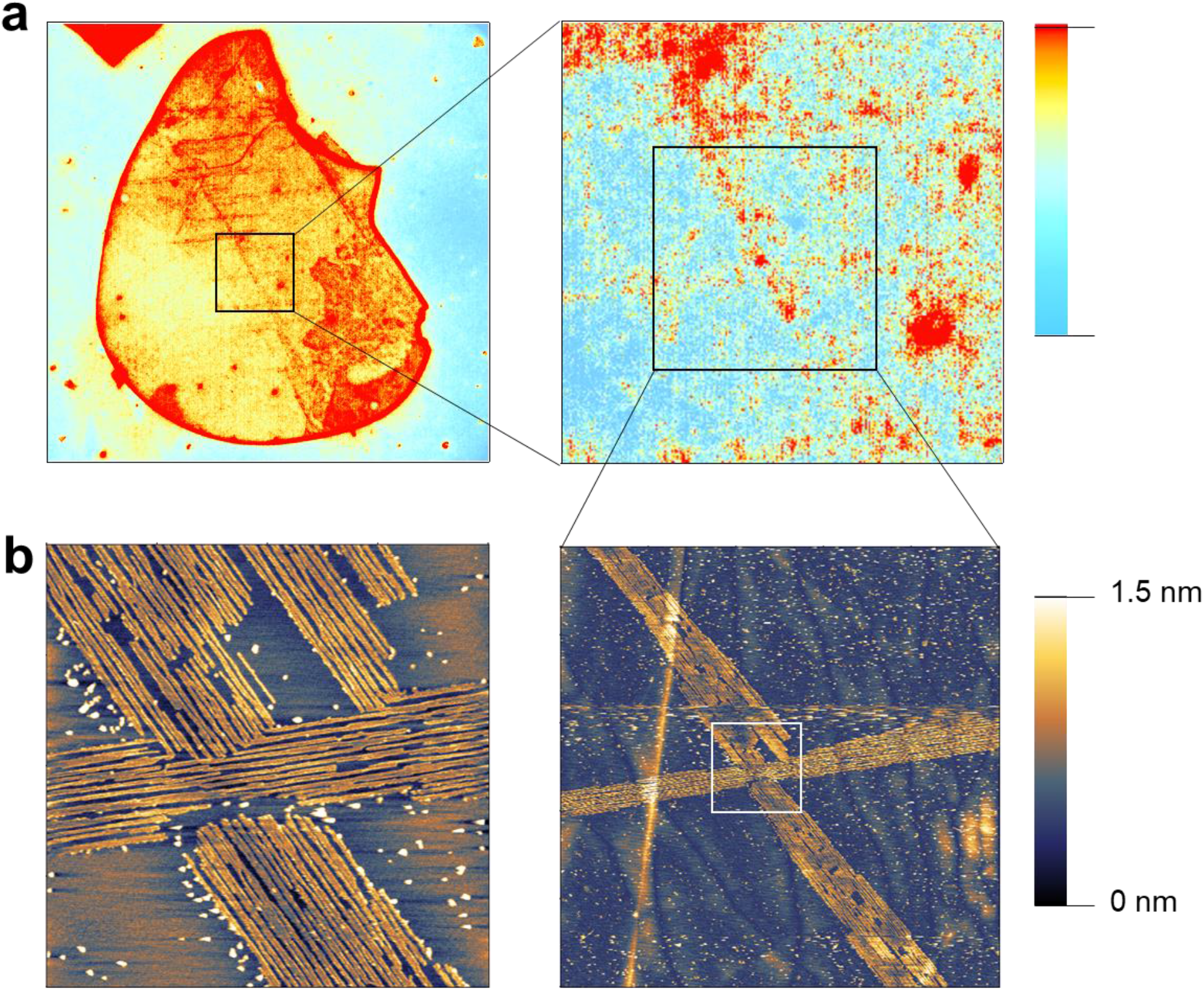
Binding of Thioflavin T to self-assembled NR7C nanostructures on the surface of boron nitride flakes, observed using fluorescence microscopy and AFM. (A) fluorescent images of an h-BN flake on a glass coverslip. (B) AFM images showing nanowire assembly on h-BN.

h-BN was chosen for the assay because of its high transparency in the visible region, making it suitable for use with the transparent substrate of the microscope slide, as required by the instrument setup (see Methods) [38]. ThT is a dye derived from benzothiazole, commonly used to detect the presence of amyloid fibrils with highly stable β-sheet structures in solution. When bound to amyloid fibrils, the dye exhibits increased fluorescence, making it an effective diagnostic tool [39]. The stacked β-sheet structures of amyloid fibrils are one of the structural and morphological characteristics shared by fibres of spider silk [40].

As shown in Figure 3a, structures resembling nanowires can be clearly observed on the surface of the h-BN flake using fluorescence microscopy. The presence of NR7C nanowires, consistent with those presented in Figure 2, was further confirmed using AFM (Figure 3b. The observation of linear domains in the fluorescent image and the agreement with the AFM image further suggest that NR7C has self-assembled on the h-BN surface into stable, highly ordered β-sheet structures with a large domain size. The linear shape of a nanowire assembled domain on the h-BN surface spans more than 50 um in length.

### Conformational characterization of spidroin NR7C by means of Raman spectroscopy

Next, we characterised the NR7C using Raman spectroscopy to seek the conformational information. Raman spectroscopy, an inherently non-invasive analytical method, detects the inelastically scattered signals emanating from molecular vibrations, intimately associated with molecular structures (Figure 4).

**Figure 4:**
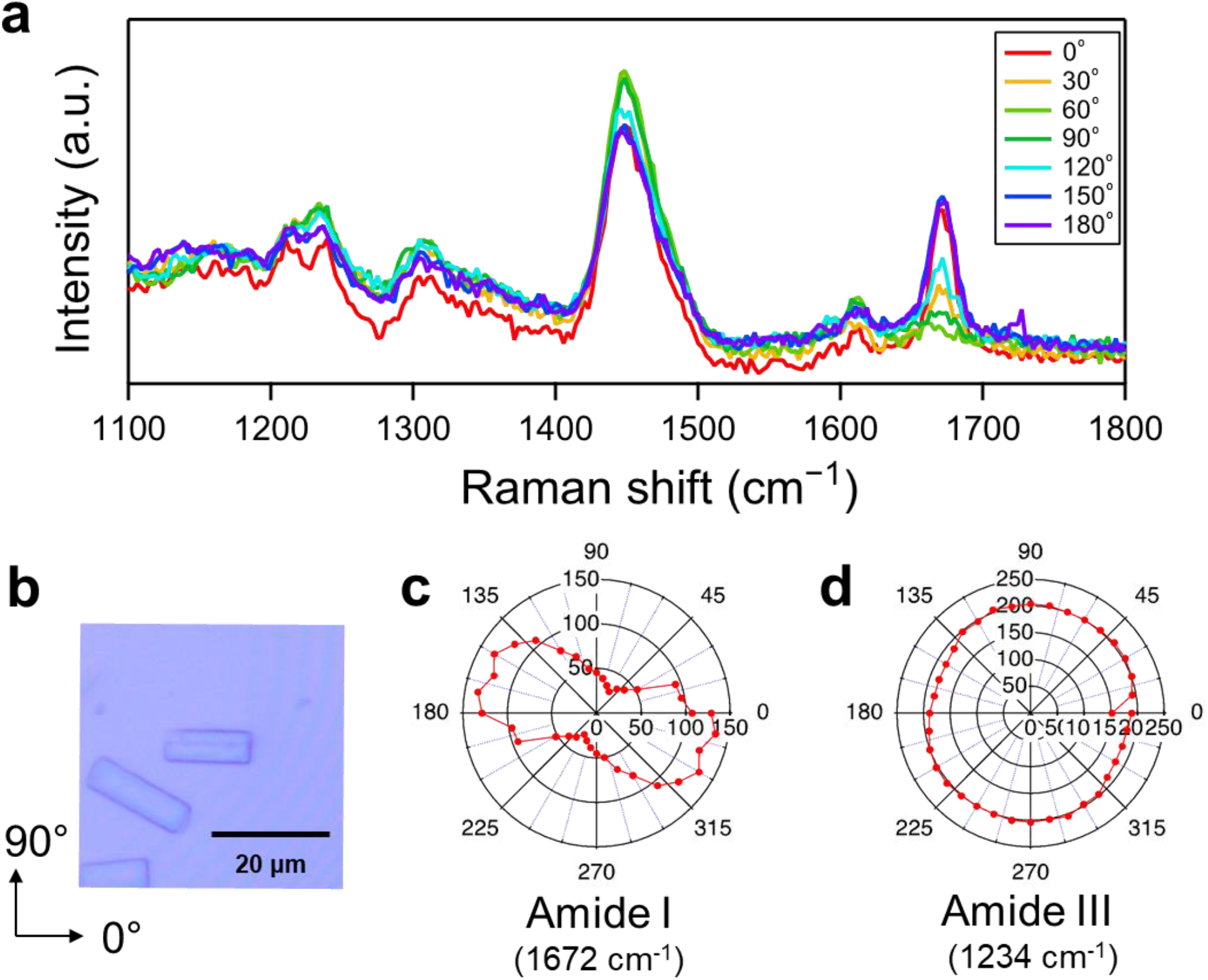
Inset A) Polarisation-resolved Raman spectra of crystallised NR7C. Inset B) Optical microscope image of NR7C crystals used in the collection of polarisation-resolved Raman spectra. Inset C) Polarisation dependence of Amide I peak intensity. Inset D) Polarisation dependence of Amide III peak intensity.

This technique facilitates the elucidation of molecular orientation and conformation in response to the incident light field. In particular, Raman spectroscopic evaluation of proteins leverages the distinctive vibrational peaks of amide bonds, which is the quintessential chemical linkages in proteins, to discern the secondary structures of proteins [41-44]. In this study, we first employed polarisation-resolved Raman spectroscopy to elucidate the secondary structures of proteins in the crystal states. This approach is powerful in assessing the molecular orientation of protein assemblies that predominantly exhibit β-sheet configurations, such as amyloid fibres and silk fibroin biomaterials [45]. The local chemical structures of the crystallised NR7C were characterised by concentrating the excitation light at the centre of the rod-shaped crystal (Figure 4b). The polarisation angle of the excitation laser was first set to parallel to the major axis of the crystal spidroin, remarked as 0 degrees. The polarisation angle was rotated every 10 degrees to investigate the polarisation dependence of Raman spectra of the crystal spidroin. The polarization-resolved Raman spectra of the NR7C crystal, acquired at varying polarization angles incremented by 30 degrees, are depicted in Figure 4a. The peaks in the Raman spectra of spidroin were assigned into vibrational modes according to Hamaguchi and Iwata [46] and are summarised in Table S1. The bands at 1674 cm^-1^ and 1231 cm^-1^ correspond to the amide I and amide III bands, respectively, and are in good agreement with the prior Raman data and the literature. The peak positions of amide I band and amide III infer the formation of β-sheet in the the dominant secondary structure of NR7C spidroin as previously reported. In order to further investigate the orientation of the spidroin proteins in the crystal, polar plots summarising the angle dependence of the intensities of the amide I and amide III peaks are displayed in Figure 4c and 4d, respectively. The intensities of the amide I band display clear anisotropic polarisation dependence, with the direction of the major axis tilted by about 30 degrees clockwise from 0 degree. This is due to the highly ordered arrangement of secondary structural elements of NR7C within the crystal lattice, suggesting the organisation of NR7C into nanofibrillar structures. On the other hand, the intensities of the amide III band exhibited clear isotropic polarisation dependence because the amide III band, which is predominantly associated with the molecular vibrations of N-H bending and C-N stretching, thereby no polarisation dependence should be observed. As can be seen from Figure 4c, the maximal Raman signal intensity for the amide I band was observed at a minor tilt along the crystal’s major axis. This observation is consistent with the reported internal orientation of spidroin within the fibrils, which is aligned at an angle along the fibril’s major axis [44].

### Tip-Enhanced Raman Spectroscopy

In order to elucidate the structural properties of self-assembly of NR7C spidroins on 2D nanomaterials, we employed a nanoscale chemical characterisation technique, called as tip-enhanced Raman spectroscopy (TERS). TERS exploits a metallic nano-tip, at which the light field is confined and enhanced owing to localised surface plasmons, resulting in generation of near-field light, to excite Raman scattering of molecules in the vicinity of the apex of the nano-tip [48-50]. Due to its high detection sensitivity and high spatial resolution, TERS has been utilised for the chemical characterisation of nanostructured samples, including 2D nanomaterials [51, 52], polymers [53], and biomolecules [54-56]. Herein, we leveraged TERS to probe local chemical and structural information on NR7C spidroin immobilised on a monolayer MoS_2_. Our TERS system consists of an inverted optical microscope with a Raman spectroscopy system coupled with AFM to precisely control the position of the metallic nano-tip. The detail of our TERS system is described in the methods section. The metallic nano-tip was brought into the vicinity of the sample and illuminated by an incident light to enhance the Raman scattering of the sample. In our TERS measurements, NR7C spidroin nanostructures and MoS_2_ layers were on the thin gold film fabricated on the glass coverslip to employ the so-called gap-mode configuration in TERS (Figure 5a).

**Figure 5:**
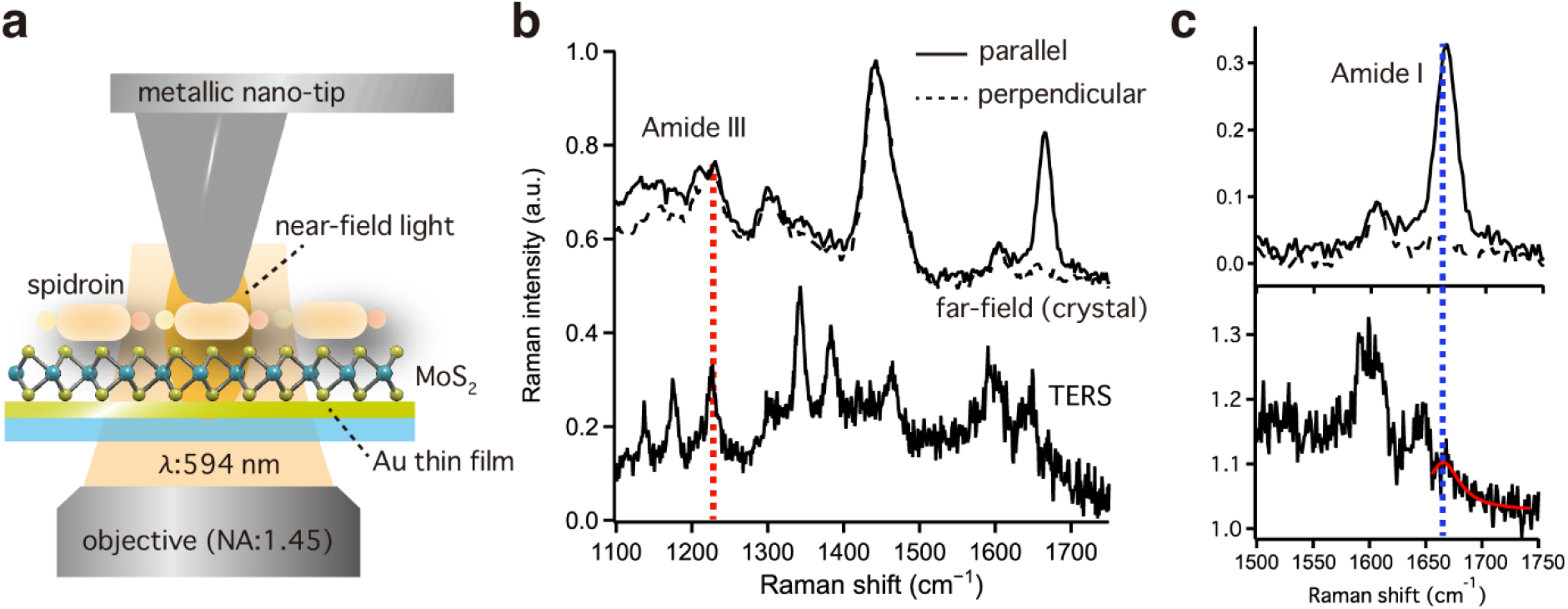
Inset A) Schematic illustration of TERS system, in which the near-field light at the apex of the metallic nano-tip excites Raman scattering from the sample. Inset B) Comparison of TERS spectrum of self-assembled NR7C mini-spidroin on MoS_2_ and polarisation-resolved Raman spectra of crystallised NR7C. The red dotted line indicates the Raman shifts of Amide III band. Insert C) Lorentzian-fitted TERS spectrum (red) of self-assembled NR7C mini-spidroin on MoS_2_ alongside far-field Raman spectra of crystallised NR7C with polarisations parallel and perpendicular to the crystal axis, detailing the region of the Amide I band. The blue dotted line indicates the Raman shift of the Amide I band in the TERS spectrum.

In the gap-mode TERS, light fields are extremely confined and enhanced at the gap between the metallic tip and the metallic substrate, which allows us to achieve single molecular Raman analysis [57, 58]. A typical optical micrograph and AFM image of the NR7C spidroin sample was shown in Figure S1 in Supporting Information to confirm the immobilisation of spidroin on monolayer MoS_2_. The typical TERS spectrum of NR7C spidroin on the MoS_2_ monolayer is shown in Figure 5(b) together with the Raman spectrum of crystal spidroin with the polarization angle of 0 (parallel) and 90 (perpendicular) degrees. Several characteristic peaks of NR7C spidroin were observed in the TERS spectrum; for example, peaks at 1138 cm^-1^, 1212 cm^-1^, 1226 cm^-1^, 1300 cm^-1^, 1345 cm^-1^, 1462 cm^-1^, and 1606 cm^-1^, which are consistent with band assignment of Raman peaks in Figure 4b. The slight variation in the frequency of vibrational modes was attributed to peak shift due to electronic interaction between molecules and metal, known as a chemical enhancement [59, 60]. The observed specific enhancement and appearance of new peaks in the TERS spectrum could be attributed to the interplay between the protein conformation and the decay nature of the enhanced optical field [61], or chemical enhancement. For identification of structural properties of spidroin, the frequencies of the amide III band and the amide I band were analysed. The frequencies of amide III band in the TERS spectrum of NR7C spidroin and the Raman spectrum of the NR7C spidroin crystal in Figure 5b were 1226 cm^-1^ and 1231 cm^-1^, both of which show the formation of β-sheet in the the dominant secondary structure of NR7C spidroin as assigned in the Raman spectrum of the crystal spidroin and previous work [44, 56]. The amide I band in the TERS spectrum of the NR7C spidroin was weak, but detectable as shown in Figure 5c, in which raw data (black) and Lorentzian-fitted data (red) of the TERS spectrum of self-assembly of spidroin were shown. The frequencies of the amide I band recorded in the TERS and Raman spectra were 1667 cm^-1^ and 1666 cm^-1^, respectively, which also indicates formation of β-sheet secondary structure in NR7C spidroin. We would like to mention here that the notably weak intensity of the amide I band in the TERS spectrum of NR7C spidroin was attributed to the polarisation dependence of the amide I band. In TERS, the polarisation of confined optical fields is parallel to the axis of the metallic nano-tip and the amide I band has strong polarisation dependence for its selection rules as proved in previous work [62] and Figure S2 in Supporting Information, where the intensity of the amide I band was diminished in z-polarised Raman spectrum of crystal spidroin samples. Hence, the weak amide I band in the TERS spectrum of NR7C spidroin on MoS_2_ monolayer suggested that the relative orientation of C=O bonds in NR7C spidroin against the polarisation of optical fields at the tip apex was perpendicular. This agrees with the fact that NR7C spidroin in crystal forms β-sheet secondary structures with the molecular axis parallel to the surface, in which the C=O bonds are primarily oriented parallelly to the surface. On the other hand, the TERS spectrum of NR7C spidroin distinctly shows the amide III band, corroborating its minimal polarization dependence, as evidenced in Figure S2, where amide III bands in the far-field Raman spectra of NR7C crystal show no polarisation dependence. The comparison between TERS spectrum of NR7C spidroin on MoS_2_ monolayer and far-field Raman spectra of both crystal and lyophilised NR7C spidroin indeed proves the formation of monolayer 2D crystal of spidroin on the MoS_2_ layer. The far-field Raman spectrum of the lyophilized spidroin sample is depicted in Figure S3. In general, Raman spectra of lyophilised form of protein exhibit peak shifts due to the weak intermolecular interactions, most probably originating from the strong hydrogen bonding at the amide bonds in the backbone as opposed to their crystal forms and random orientations of proteins in the focal volume [63]. The amide I band and amide III band in the lyophilised NR7C appears at 1671 cm^-1^ and 1242 cm^-1^, respectively. These frequencies exhibit slight variations when compared to the Raman spectra of the crystalline and self-assembly of NR7C. In other word, the good agreement of amide I and amide III band frequencies as well as those of other hands in the TERS spectra of self-assembly of NR7C with those found in the Raman spectrum of its crystalline form leads to the conclusion that the 2D crystal of NR7C spidroin proteins characterised by a highly ordered β-sheets conformation has been formed on the MoS_2_ layer owing to the strong intermolecular interactions.

## Conclusions

This study reports the first demonstration of self-assembly of a recombinant spider silk protein across 2D material surfaces. Self-assembly into ordered, long-range nanostructures was confirmed using AFM and fluorescence microscopy. The structural conformation of NR7C crystal was further characterised using Raman spectroscopy, revealing a highly stable, predominantly β-sheet structure. Finally, conformational properties of self-assembly of NR7C on a monolayer MoS_2_ were unveiled at single protein level by TERS measurement. The formation of 2D crystal layer of NR7C on the monolayer MoS_2_ was successfully demonstrated. Our results indicate that recombinant proteins based on spider silks possess significant potential for use as scaffolds in molecular biosensing. Future studies will examine surface self-assembly of recombinant spider silk proteins containing repetitive regions of varying sizes. In addition, spidroin solutions will be prepared in a collection of buffer systems based on the key factors known to influence fibre formation, namely pH and ion gradients. This will enable the development of a series of rules for influencing fibre self-assembly at the material surface, such as coverage and directionality. Finally, the biosensing ability of any candidate spidroin scaffold must be examined via the immobilisation of probe molecules.

## Materials and methods

### Preparation of NR7C spidroin samples

Recombinant spidroin NR7C was expressed, purified and lyophilised as described in Finnigan et al [23]. Lyophilised spidroin powder was stored at -20 °C prior to use and was resuspended in DI water as required. Resuspended spidroin solutions were filtered using hydrophilic PTFE filters (0.20 μm pore size) to remove undissolved aggregates. Stock spidroin solutions were prepared through serial dilution in DI water. Concentrations of resuspended spidroin samples were determined in triplicate using a UV-vis spectrophotometer (NanoPhotometer, Implen). The molecular weight and A_280_ extinction coefficient of recombinant spidroin NR7C were calculated using the ExPaSy ProtParam tool [36]. The amino acid sequence of NR7C is presented in Table S2.

### Preparation of samples for Atomic Force Microscopy

For observation of spidroin self-assembly on 2D material substrates (graphite, MoS_2_ and boron nitride), flakes of each material were transferred onto a silicon wafer containing a 300 nm layer of silicon dioxide (SUMCO, Japan) via the mechanical exfoliation method detailed in Novoselov et al. [64]. Substrate flakes were annealed to the SiO_2_ wafers (200 °C, 30 minutes). 2D material substrate samples were incubated with 50 μl of spidroin solution at room temperature for one hour. Following incubation, the solution was removed using nitrogen gas and dried in a vacuum desiccator. Sample preparation is summarised in Figure S4.

### Atomic Force Microscopy measurement

Spidroin self-assembly on 2D material surfaces was characterised using a commercially available AFM (Agilent Technologies 5500 Scanning Probe Microscope, Agilent Technologies). AFM measurement was performed in ambient condition and equipped with a silicon cantilever (OMCL-AC160TS, Olympus). Samples were characterised using a resonance frequency of 300 kHz and a spring constant of 26 N/m.

### AFM image processing

Image processing software Gwyddion (Czech Metrology Institute, CZ) [65].

### ThT fluorescence measurement

Microscope coverslips were spin-coated with Poly(methyl methacrylate). Flakes of boron nitride were transferred onto the coated coverslips using the mechanical exfoliation method described in Novoselov et al. [64]. The substrate was wetted using 100 μl DI water prior to the addition of 100 μl spidroin solution. Samples were incubated at room temperature for one hour. Post-incubation, the spidroin solution was exchanged with DI water at the substrate surface. Thioflavin T (ThT) solution was added to the substrate surface to a final concentration of 50 nM. Samples were then incubated for an additional hour. Following incubation with ThT, samples were blown dry with nitrogen gas and dried under vacuum.

To examine ThT fluorescence, the samples were placed on an inverted fluorescence microscope (IX73P2F, Olympus) with a fluorescence filter cube (emission 420-460 nm; excitation >515 nm). The microscope was equipped with an oil-immersion lens (NA = 1.4) and a CMOS camera (Neo sCMOS/Solis, Andor). A mercury lamp (870 μW) was used to excite the bound ThT, and a series of images were recorded continuously to investigate the interaction between ThT and the self-assembled spidroin structures on the h-BN flakes.

### Crystallisation of NR7C spidroin

Crystallisation studies were conducted using the sitting-drop method. NR7C spidroin solution was prepared to a final concentration of 1 mg/ml in 10 mM potassium phosphate buffer, pH 7. Spidroin solution was applied to a 96-well screening plate containing 10% (w/v) NaCl solution within each reservoir. Plates were incubated at 25 °C. Protein crystals appeared after 4 days of incubation.

### Polarised Raman spectroscopy of crystallised spidroin

Raman spectra of crystallised spidroin were collected using a home-made optical system coupled with a spectrometer (Isoplane 320, Princeton Instruments) and a peltier-cooled CCD camera (Pixis 400, Princeton Instruments). A continuous wave laser with a wavelength of 532 nm was used for Raman excitation, with a power density of 77 μw/μm^2^. Samples of crystallised spidroin were monitored and irradiated using a dry objective lens (x 40, N.A. = 0.7), over an exposure time of 100 seconds. Laser polarisation was achieved using a Glan-Thompson prism. The laser polarisation angle was controlled using a Fresnel rhomb 1/2 waveplate. The polarization dependence of the spectrometer was countered by placing a 1/4 waveplate in front of the spectrophotometer. The setup is summarised in Figure S5.

### Raman spectroscopy of lyophilised spidroin

Raman spectra of lyophilised spidroin powder were collected using a different Raman spectroscopic system mentioned above. A He-Ne laser (Thorlabs Inc., HNL150LN) was used for excitation with a power of 7.18 × 10^3^ μw/μm^2^. The excitation wavelength was 632.8 nm. Raman-scattered light was recorded using a spectrophotometer (Acton SP 2358-T, Princeton Instruments) coupled to a CCD camera (Pixis 100, Princeton Instruments). Samples were prepared by sandwiching lyophilised spidroin powder between a silicon wafer and a glass coverslip. Spectra were obtained using a 100x objective lens (NA=0.85), over an exposure time of 100 seconds.

### Tip-enhanced Raman spectroscopy

All TERS measurements were performed using a bottom-illumination AFM-based TERS system that incorporates an inverted optical microscope and an AFM system (MFP-3D, Oxford Instruments) [66, 67], a spectrometer (IsoPlane 320, Teledyne Princeton Instruments) and a peltier-cooled CCD camera (PIXIS 100, Teledyne Princeton Instruments) were coupled with the inverted optical microscope. Unlike typical TERS measurements for small molecules or thin samples, such as benzenethiol molecules and 2D materials, our sample for TERS measurements consists of spidroin and monolayer MoS_2_, which produce the relatively large gap distance of at least 2 nm between the metallic tip and the metallic substrate. In such a case, the plasmonic resonance wavelength should appear in the relatively shorter wavelength region around 600 nm. For this reason, a continuous wave laser with a wavelength of 594 nm (Mambo, Cobolt) was employed as an incident light source. The power density of the incident laser at the sample plane was set to less than 400 μw/μm^2^ to avoid photodamage of fragile biological samples [62]. A commercially available oil immersion objective (×100, N.A. 1.45) was used for both the excitation and collection of TERS signals. The backscattered optical signal was first filtered by a sharp-edge long-pass filter, and only the Raman scattering signal was guided into the spectrometer. The TERS system was enclosed by a simple enclosure to avoid the possible thermal drift of AFM. The contact mode operation of AFM was employed for TERS measurement to maximise the plasmonic enhancement of Raman scattering.

TERS spectral data were analysed by Igor software. TERS spectra represent the pure intensity of enhanced Raman signal, which is obtained after subtracting a Raman spectrum measured when a metallic tip is far away from a sample from the one measured when a metallic tip is near a sample.

Metallic probe tips were fabricated by physical vapour deposition processes. Commercially available AFM silicon cantilever tips were pre-cleaned by UV-Ozone cleaning, and a 40-nm-thick silver layer was subsequently thermally evaporated and deposited on the tips.

A thin gold film substrate for TERS measurements was fabricated through the thermal evaporation of Ge and Au. A 2-nm-thick Ge buffer layer was first pre-coated on a cleaned glass coverslip, and Au was thermally evaporated onto the substrate to form an ultrasmooth gold layer with a thickness of 10 nm. Owing to the Ge buffer layer, the surface roughness of the thin gold film was as low as 1.0 nm, which avoids any hot spots on the Au surface [68]. Thus, the Au thin film should not bring Raman signal enhancement alone.

Next, single-layer of MoS_2_ was grown on a Si wafer using a chemical vapor deposition method. The MoS_2_ was then transferred onto a glass coverslip coated with Au/Ge using a stamp of polydimethylsiloxane (PDMS) coated by a polystyrene (PS) support film. After transferring the MoS_2_ and PS support film, the glass coverslip was soaked in a toluene solution at 90°C for 30 minutes, and this process was repeated three times to remove the PS film. The sample with transferred MoS_2_ was finally rinsed with toluene at room temperature and dried using nitrogen gas. Typical optical and AFM image of the sample was shown in Figure S1 in Supporting Information.

It should be noted that the enhanced Raman scattering of the sample could be obtained only when the metallic tip is in the vicinity of the sample as confirmed by Figure S6 (a) in the Supporting Information. The TERS spectrum of a flat Au thin film, on which no molecule exists was also recorded using the metallic tip employed for successful TERS measurement to confirm contamination of the TERS signal did not occur (Figure S6 (b)).

## Acknowledgements

We are grateful to W. Finnigan for supplying recombinant spidroin NR7C protein.

D.R.W. was supported by a JSPS postdoctoral fellowship (PE21770).

R.K. acknowledged the financial support from JSPS KAKENHI grant no. JP22K14650, JST ACT-X grant no. JPMJAX21B4, Tateishi Science and Technology Foundation, and Murata Science and Education Foundation.

R.K., A.O., and T.Y also acknowledge the support from the project on the Promotion of Regional Industries and Universities by the Cabinet Office.

E.T. was funded via Engineering and Physical Sciences Research Council (EPSRC), Grant Ref: EP/X015408/1.

Y.H. was supported by JSPS KAKENHI Grant Numbers 25706012 and 16H05973, Grant-in-Aid for Transformative Research Areas 22H05408, and the Precise Measurement Technology Promotion Foundation (PMTP-F).

## Author contributions

D.R.W., R.K. and Y.H. wrote the manuscript. D.R.W. performed atomic force microscopy and fluorescence microscopy experiments. R.K. performed TERS experiments. A. O. assisted with TERS experiments. R. T. performed atomic force microscopy experiments. K. M. performed angle-resolved Raman and protein crystallisation studies. H. M. assisted with fluorescence microscopy experiments. Y. Y. performed Raman experiments on lyophilised protein. T. Y., E. T., and Y. H. designed experiments and analysed data. All authors contributed to the proofing of the manuscript. The present manuscript was submitted with agreement from all authors.

