## Supplementary Figures S1-S6, Supplementary Tables S1-S2 for "Synthetic spider silk forming highly-aligned nanoarchitectures on 2D-material surfaces"

**Supplementary information**

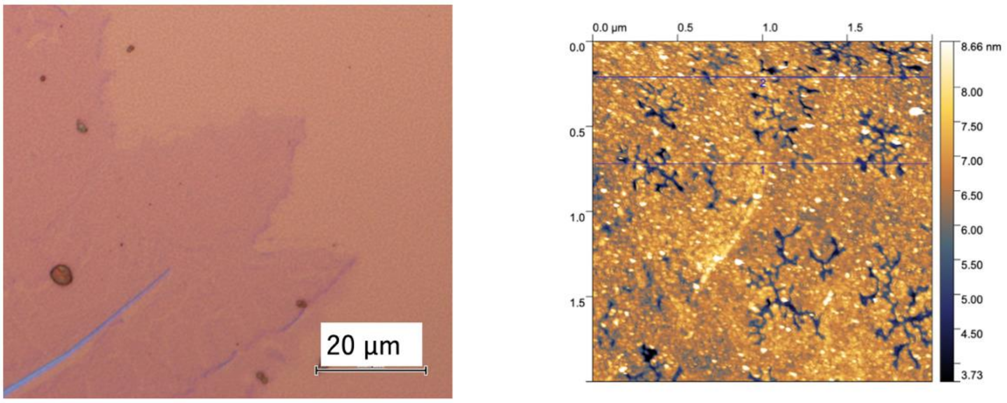


***Figure S1:*** *Inset A) Optical micrograph of NR7C mini-spidroin on the monolayer $MoS_2$ showing representative areas for TERS measurements. Inset B) Representative AFM image of the NR7C mini-spidroin sample for TERS measurement.*

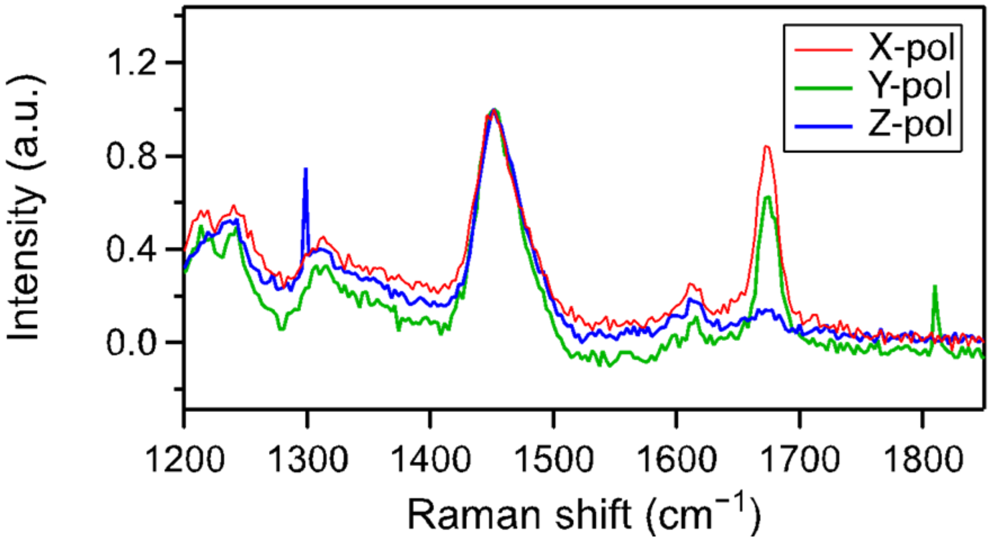


***Figure S2:*** *Polarisation dependence of Raman spectra of crystal form of NR7C spidroins. (normalised by the peak near 1450 cm$^{-1}$).*

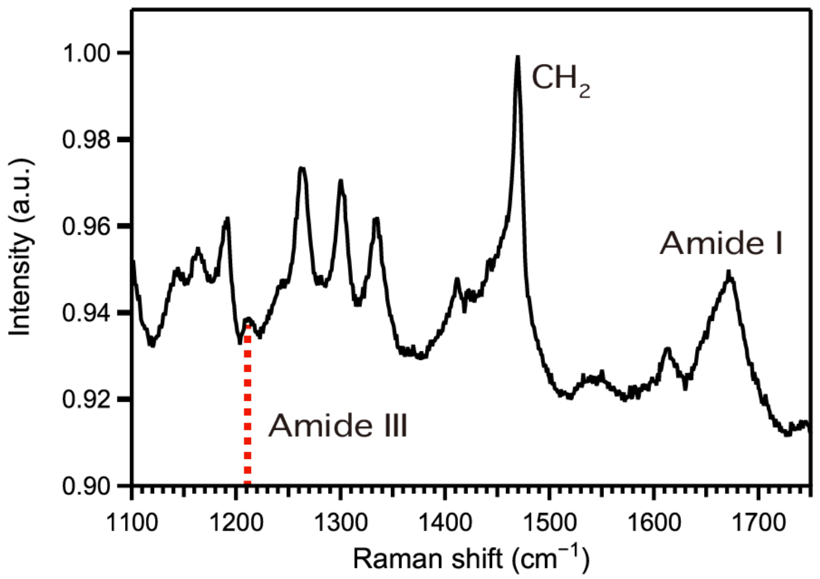


***Figure S3:*** *Far-field Raman spectrum of lyophilised NR7C.*

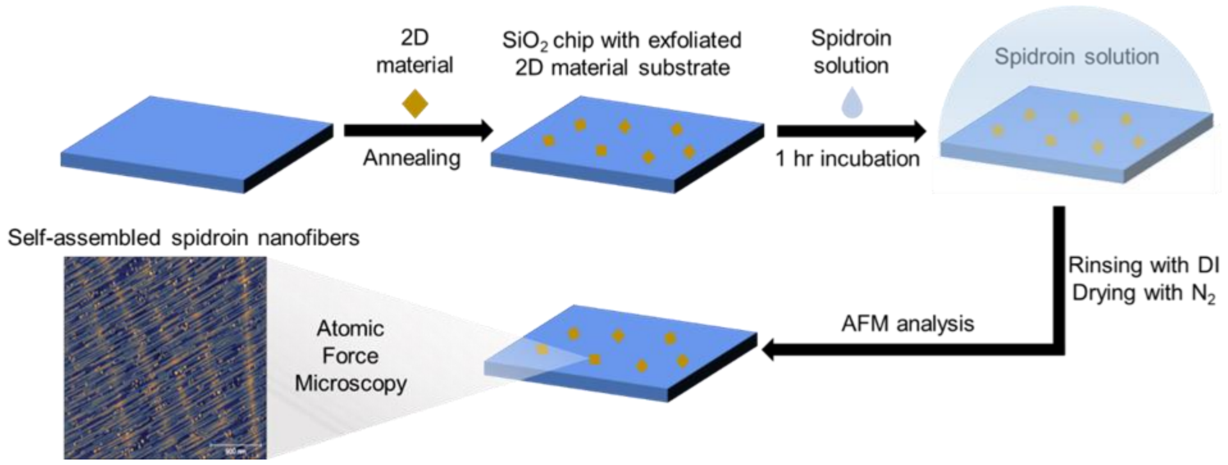


***Figure S4:*** *Summary of sample preparation methodology for AFM observation of self-assembled spidroin nanostructures on 2D material substrates.*

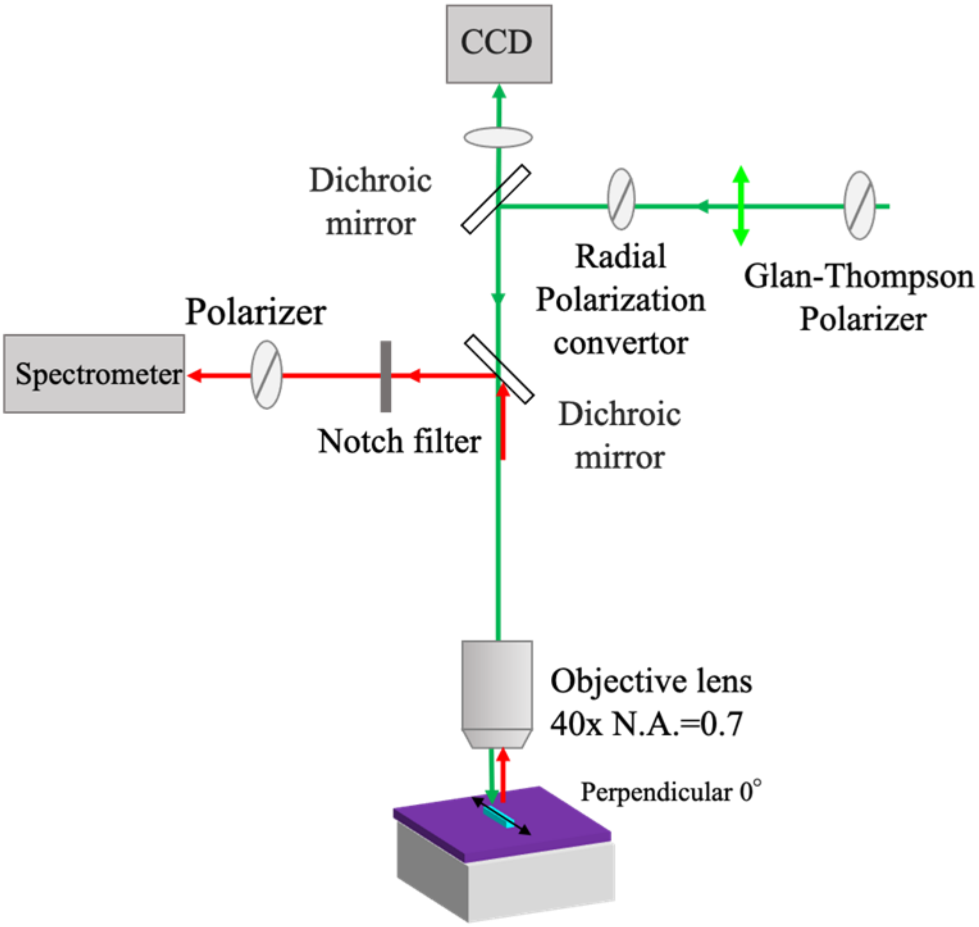


***Figure S5***: *Schematic of instrument setup used for polarised Raman spectroscopy.*

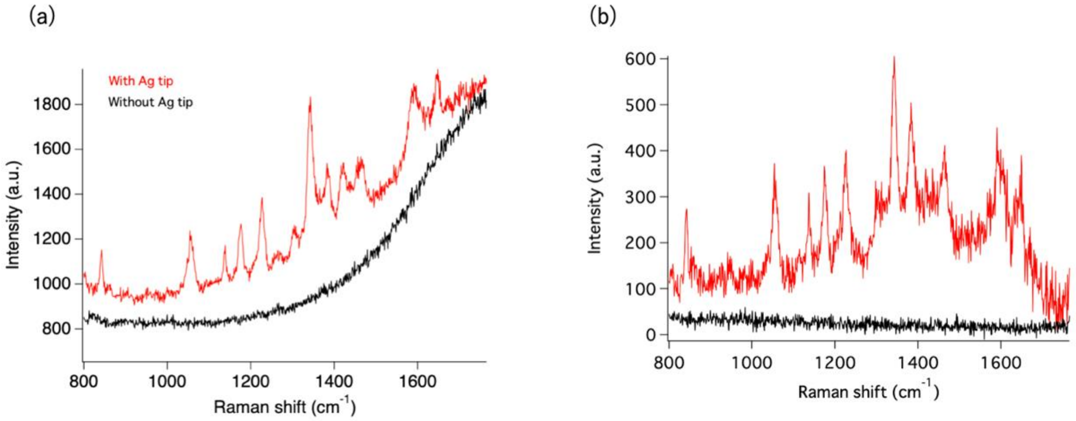


***Figure S6:*** *Inset A) Comparison of TERS spectra of NR7C mini-spidroin on $MoS_2$ when the tip is in contact with (red) and away from (black) the sample. Inset B) Comparison of TERS spectra of NR7C mini-spidroin on $MoS_2$ (red) and the flat Au thin film (black)*

***Table S1***: *Summary of peak assignments from crystallised NR7C angle-resolved Raman spectra. All peaks assigned according to Hamaguchi and Iwata [1].*

| Peak number | Peak position ($cm^{-1}$) | Peak assignment |
|---|---|---|
| **1** | 1138 | Skeletal<br>C-C(trans) stretch. |
| **2** | 1212 | $C-C_6H_5$<br>(phenyl ring) stretch. |
| **3** | 1226 | - |
| **4** | 1233 | Amide III |
| **5** | 1300 | $CH_2$ twist. |
| **6** | 1304 | $CH_2$ wag |
| **7** | 1345 | CH deform. |
| **8** | 1448 | $CH_2$ bend |
| **9** | 1462 | $CH_3$ deg. deform. |
| **10** | 1606 | Phe |
| **11** | 1613 | Tyrosine, Tryptophan |
| **12** | 1671 | Amide I |

***Table S2:*** *Protein sequence for recombinant spidroin, NR7C.*

**>NR7C**
**MHHHHHHSSGVDLGTENLYFQSMALGQANTPWSSKENADAFIGAFMNAASQSGAFSSDQIDDMSVISNT**
**LMAAMDNMGGRITQSKLQALDMAFASSVAEIAVADGQNVGAATNAISDALRSAFYQTTGVVNNQFITGIS**
**SLIGMFAQVSGNEVAGGGAGQGGQGGYGRGGYGQGGAGQGGAGAAAAAAAAGGAGQGGQGGYGQG**
**GYGQGGAGQGGAAAAAAAAAGGAGQGGYGRGGAGQGGAGGSGPGQIYYGPQSVAAPAAAAASALAA**
**PATSARISSHASALLSNGPTNPASISNVISNAVSQISSSNPGASACDVLVQALLELVTALLTIIGSSNIGSVNYDSS**
**GQYAQVVTQSVQNAFAGS***
